# Volume growth in animal cells is cell cycle dependent and shows additive fluctuations

**DOI:** 10.1101/2021.06.03.446986

**Authors:** Clotilde Cadart, Matthieu Piel, Marco Cosentino Lagomarsino

## Abstract

The way proliferating animal cells coordinate the growth of their mass, volume, and other relevant size parameters is a long-standing question in biology. Studies focusing on cell mass have identified patterns of mass growth as a function of time and cell cycle phase, but little is known about volume growth. To address this question, we improved our fluorescence exclusion method of volume measurement (FXm) and obtained 1700 single-cell volume growth trajectories of HeLa cells. We find that, during most of the cell cycle, volume growth is close to exponential and proceeds at a higher rate in S-G2 than in G1. Comparing the data with a mathematical model, we establish that the cell-to-cell variability in volume growth arises from constant-amplitude fluctuations in volume steps, rather than fluctuations of the underlying specific growth rate. We hypothesize that such “additive noise” could emerge from the processes that regulate volume adaptation to biophysical cues, such as tension or osmotic pressure.

## Introduction

The regulation of animal cell growth is a central question in cell biology^1–3^, but our knowledge is limited by the lack of methods to reliably measure cellular growth at the single-cell level. In the last decade, several sophisticated approaches measuring buoyant mass^4,5^, dry mass^6–9^ and volume^10,11^ have produced new data revealing unexpected features at several levels. In particular, in contrast to what has been observed in unicellular organisms such as *S. pombe*^12,13^, *S. cerevisiae*^14^ or *E. coli*^15^, growth patterns of single animal cells *in vitro* cannot easily be associated to a simple growth mode, such as mono-exponential, linear or bi-linear. Instead, single animal cells show complex growth patterns that remain poorly understood to date. Note that to avoid ambiguity we hereon call “growth speed” the time-derivative of mass or volume (e.g., 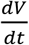) and “mass- or volume-specific growth rate”, the growth speed divided by mass or volume (e.g., 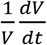).

So far, studies on single animal cell growth have focused on patterns observed at timescales ranging from several hours to a cell cycle. For HeLa cells, growth was reported to couple with cell size, thus contributing to cell size homeostasis^10,16^. These cells were reported to grow, while in G1, at a faster-than-average volume-specific growth rate if they were born smaller than average^10^. This finding was paralleled by the observation that inhibition of cell cycle progression or growth pathways has antagonistic effects on mass-specific growth rate or cell cycle progression respectively^17^. A second type of growth pattern was identified in studies measuring cell mass, and showed an association between mass-specific growth rate and cell cycle progression or cell age. One study showed that in L1210 cells that undergo polyploidization, mass-specific growth rate follows a bell-shaped dependency on mass over the course of each cell cycle, independently of the increasing mass of the cell as ploidy increases^18^. This suggests that the bell-shape pattern of mass-specific growth rate is a function of cell cycle progression, not mass itself. Two other studies reported that the mass growth speed of several cell types displayed periodic oscillations. Although the characteristics of the oscillations identified differ in the two studies, these results show that mass growth follows an oscillatory pattern that depends on time since birth^19^ or time until division^9^.

At shorter timescales (one hour or less) single adherent cells display cell volume^10^ and cell mass^5,9^ growth trajectories that vary in time and across cells. These fluctuations have not yet been analyzed in detail, and their origin remains poorly understood. Mass or volume changes in a given time interval are the combined consequence of biosynthesis (via transcription and translation) and mechanisms that import or export mass (import of molecules via endocytosis/exocytosis)^20^ or volume (import of water via osmotic balance, hydrostatic pressure, and membrane turnover)^1^. Variability in growth can thus result from variability in either or both categories of mechanisms. While seminal studies have revealed the origins of “noise” in transcription in animal cells^21,22^, the processes leading to noisy mass and volume growth in animal cells still need exploration.

Crucially, with most of the above-cited studies focusing on cell mass, volume growth remains poorly characterized, although it is clear that it follows independent patterns from cell mass throughout the cell cycle^11,23,24^. Regarding cell volume, open questions remain regarding both the identification of a mean growth mode (e.g. linear or mono-exponential), and on the determination of the fluctuations around this trend. While the study of mean trends is possible with medium size datasets (hundreds of cells), the study of fluctuations in growth requires much larger datasets (thousands of cells), which remain difficult to generate for animal cells. In a previous study^10^, we dynamically measured the volume of single adherent animal cells using a fluorescence exclusion technique (FXm)^25^. Our results indicated that, on average, volume growth speed increases with cell volume but the data obtained were not sufficient for deeper analysis. Here, we report an improvement of the FXm method that produces higher throughput in volume readouts. We obtained a data set of around 1700 single-cell volume curves of HeLa cells, combined with the tracking of key cell cycle transitions (birth, G1/S, and mitosis). The data show that volume-specific growth rate depends on both cell cycle phase and cell volume. Our unprecedented statistical resolution also allows us to investigate the variability in volume growth and to show that it arises from constant-amplitude additive fluctuations of growth speed rather than from fluctuations of the specific growth rate.

## Results

### An improved FXm method produces high-throughput dynamic measurements of single-cell volume

To obtain high-throughput measurements of volume growth of single cells, we improved the fluorescence-exclusion based measurement (FXm) of cell volume we previously developed^11,25^. Briefly, this method relies on seeding cells in chambers of known height in the presence of a fluorescent probe (10 kDa dextran) that does not enter or harm the cell (Figure 1 a-b). Since the cell excludes the dye, the measured fluorescence in the area containing a cell is negatively proportional to the volume of that cell.

**Figure 1.**
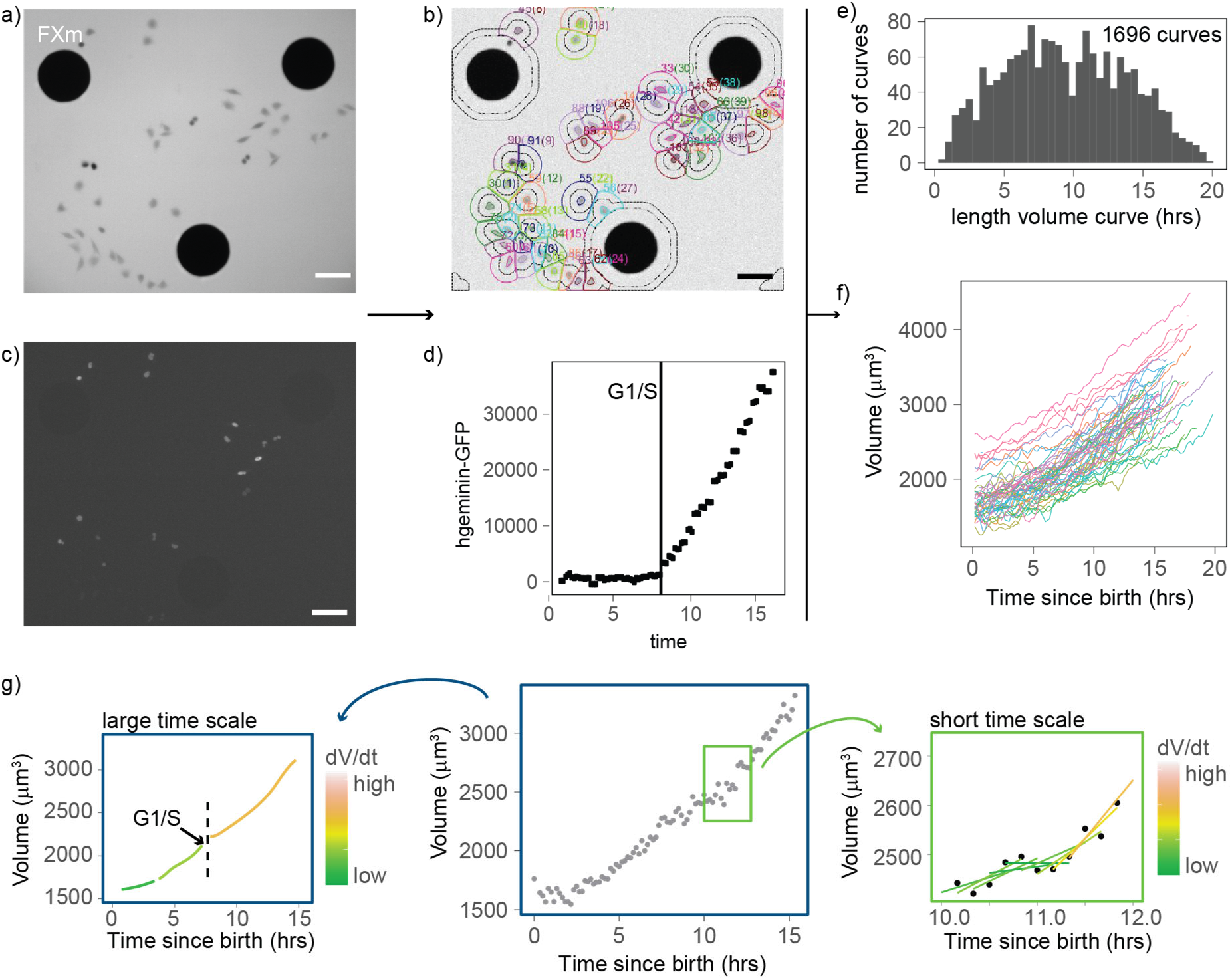
An improved FXm method produces high-throughput dynamic measurements of single-cell volume curves. a) Representative image of a field acquired in an FXm device. The FXm chamber contains a fluorescent dye (Dextran-Alexa-645) that is excluded from the cells and the pillars of the chamber. Hence, the pillars appear in black (large circles), cells are grey, and the background is bright. b) FXm images are automatically segmented using custom Matlab® software, then volume is calculated for each time point. c) Same field as in a) but imaging hgemin-GFP, a cell cycle marker expressed in the nuclei of the cells. (b,c,a: scale bar indicates 100 μm). d) Representative curve of hgeminin-GFP over time in the cell cycle for a single cell. The change of slope in the signal marks the G1/S transition. e) Histogram of the duration of each single-cell volume curve measured. We obtained a total of 1696 curves. f) 72 representative single-cell volume curves from birth to mitosis. g) Representative single-cell volume curve (middle panel) showing trends at time scales of several hours (left panel) and around one hour (right panel). Volume growth speed (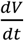) was defined from these plots as the time derivative of volume *vs* time.

We combined these volume measurements with cell cycle phase analysis using the hgeminin-GFP part of the FUCCI system^26,27^ the expression of which marks S phase entry (Figure 1c). We performed four independent 24-hour-long experiments in which we imaged thousands of cells growing asynchronously in the FXm chambers (Figure 1 – supplement 1). To extract cell volume through time for each individual cell, we developed an automated cell tracking algorithm (Figure 1b) and we verified that the segmentation and lineage tracing (recording of mitotic events) were accurate by manual inspection of each single cell trace. Birth was defined as the onset of cytokinesis, the G1/S transition was defined as the onset of increase in hgeminin-GFP intensity (Figure 1c) and mitosis was defined as the onset of the mitotic volume overshoot^10,11,24^. Using this approach, we obtained 1696 verified single-cell volume trajectories that contained all available cell cycle information (time of birth, time of G1/S and/or time of entry into mitosis, Figure 1e-f). This high-quality, high-throughput measurement of animal cell volume was used to analyze the patterns and regulation of cell volume growth with unprecedented statistical resolution.

### Volume growth is close to exponential for a wide range of volumes

First, we considered the growth mode of cells - a central question to understanding both cell growth and size homeostasis^28,29^. Typical limit cases are linear or exponential growth. As previously reported for adherent cell types^9,10^, single-cell trajectories show highly variable behavior (Figure 1g), making it difficult to associate them with any simple behavior. We turned to an alternative method based on population averages. We reasoned that, in the case of an average simple exponential growth model, growth speed should increase, on average, linearly with volume and the slope *α* followed by 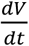 vs *V* (formally the trend of a conditional average) can be used to define an average volume-specific growth rate^30^. All four experiments consistently showed that, for volumes higher than 1800 μm^3^ (and lower than 4000 μm^3^), growth speed increases on average linearly with volume, with a slope *α*. that was very close to the (unconditional) average of 〈*1/V dV/dt*〉 (Figure 2a). The agreement of these two different estimates supports the idea that average exponential growth describes these data well. All four experiments were also very similar with values of *α*. ranging from 0.038 to 0.047 h^-1^. Thus, we conclude that volume growth is faster than linear, and on average close to exponential in this range of cell sizes. Cells with volumes below 1800 μm^3^ did not follow the same trend, likely due to a different pattern of growth early in the cell cycle (see below).

**Figure 2:**
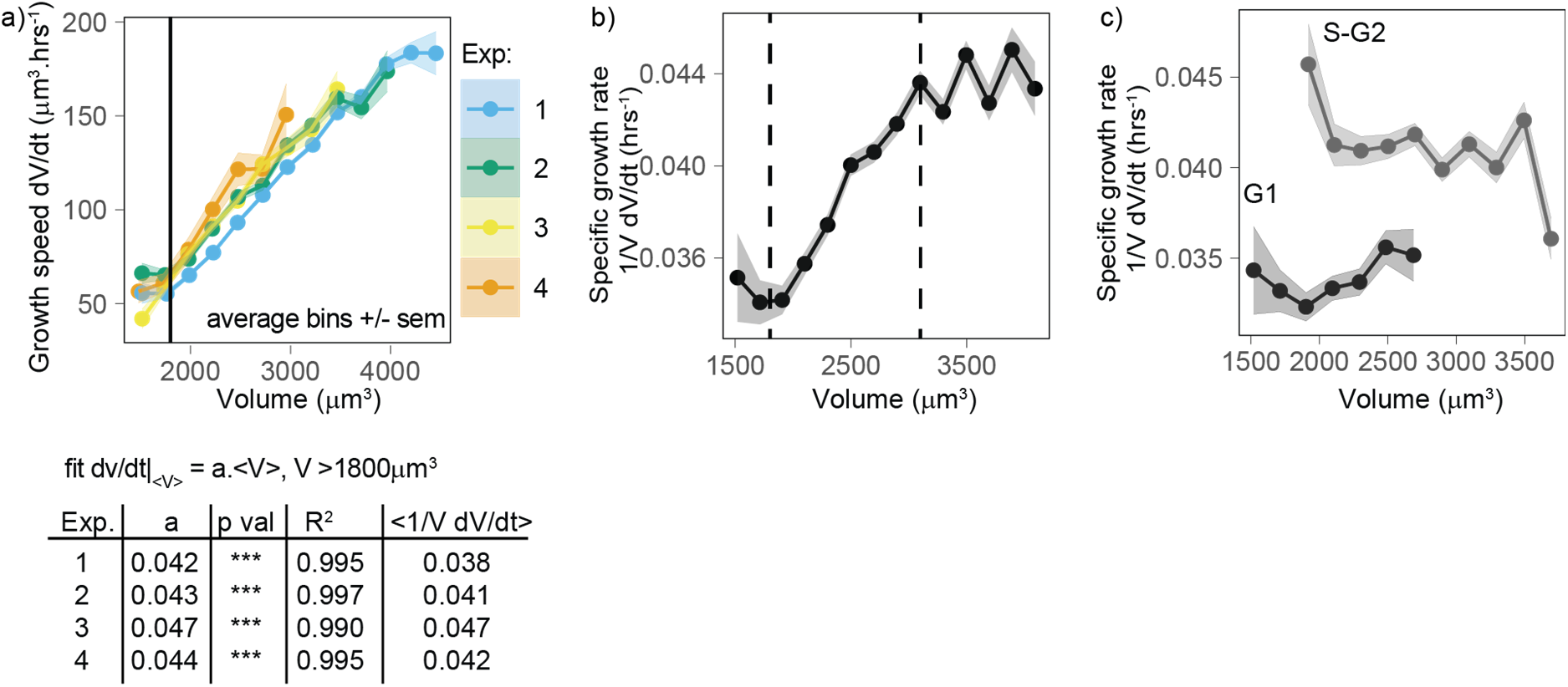
Volume growth is close to exponential on average for a wide range of volumes, with a higher specific rate in S-G2 than in G1. a) Top panel: Growth speed 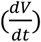 (in units volume/time) as a function of volume for each experiment (points are average bins of width = 250μm^3^, the ribbon represents the standard error on the mean for each bin, N=4, n> 25 different cells in each bin). Bottom panel: Table showing the results of the linear fit of growth speed as a function of volume for each experiment. Derivatives are computed on 50 min windows. b) Specific volume growth rate, defined as the binned average of 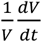 at fixed volume *V*, plotted as a function of volume (average bins of width = 200μm^3^ ± standard error on the mean). Vertical dashed lines indicate the range where growth rate increases linearly with volume, between 1800 and 3100μm^3^. (dots are average bins of width = 200μm^3^, the ribbon represents the standard error on the mean for each bin, N=4, n>100 different cells per bin). c) Same as b) but grouped by cell-cycle stage (G1 *vs* S-G2).

### Specific volume growth rate depends on cell-cycle progression

When we considered in more detail an estimated volume-specific growth rate, defined by the conditional average of 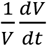 *vs* volume *V* (which has units 1/time), as a function of volume, we observed a slight but significant increase with volume for cells between 1800 and 3100 μm^3^ (Figure 2b). Outside this range, the trend is more complex but the robustness of the observed behavior may be limited due to the lower number of observations at these extreme sizes. We hypothesized that one potential cause of increase in volume-specific growth rate during the cell cycle could be a cell-cycle stage dependency, which we could test using data on the transition from G1 to S-G2 phase. To address this question, we repeated the plot of volume-specific growth rate as a function of volume, also grouping the data by cell cycle phase (G1 *vs* S-G2). This analysis shows that estimated volume-specific growth rate is nearly constant with volume for a given phase, and growth rate in S-G2 is about 15% higher than in G1 (Figure 2c). These results show that progression from G1 into S/G2 is accompanied by an increase in the volume-specific growth rate.

### Newborn cells show a distinct pattern of volume growth

The observation that small cells show a different growth behavior than the rest of the cells (Figure 2a-b) prompted us to test whether growing cells could follow different patterns early in the cell cycle. We examined volume growth speed 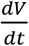 as a function of time from birth for the three experiments that had more than 80 cells (to ensure that we had enough statistical power, even when analyzing each experiment separately). In all three experiments, volume growth speed showed a fast increase during the first 1.3 hours after birth and increased more slowly and steadily after that point in time (Figure 3a). This suggests that during the initial 1.3 hours after birth, cells follow patterns different from those followed during the rest of the cell cycle. To gain more insight into the details of volume growth during this initial period, we pooled the three experiments together and grouped cells by their volume at birth. The largest cells at birth started their cell cycle with a negative growth speed (meaning that they were losing volume) during the first hour after birth. Small, intermediate, and large cells ultimately converged on the same growth speed at 1.3 hours after birth (Figure 3b). We note that in our data, birth is defined as the first time point of cytokinesis onset (a process which is then typically completed within 20-30 minutes^10^), thus the 1.3 hour period comprises the end of cytokinesis as well as early G1 phase. The fact that volume follows a distinct pattern early in the cell cycle suggests a different mechanism of volume growth regulation as cells re-enter interphase.

**Figure 3:**
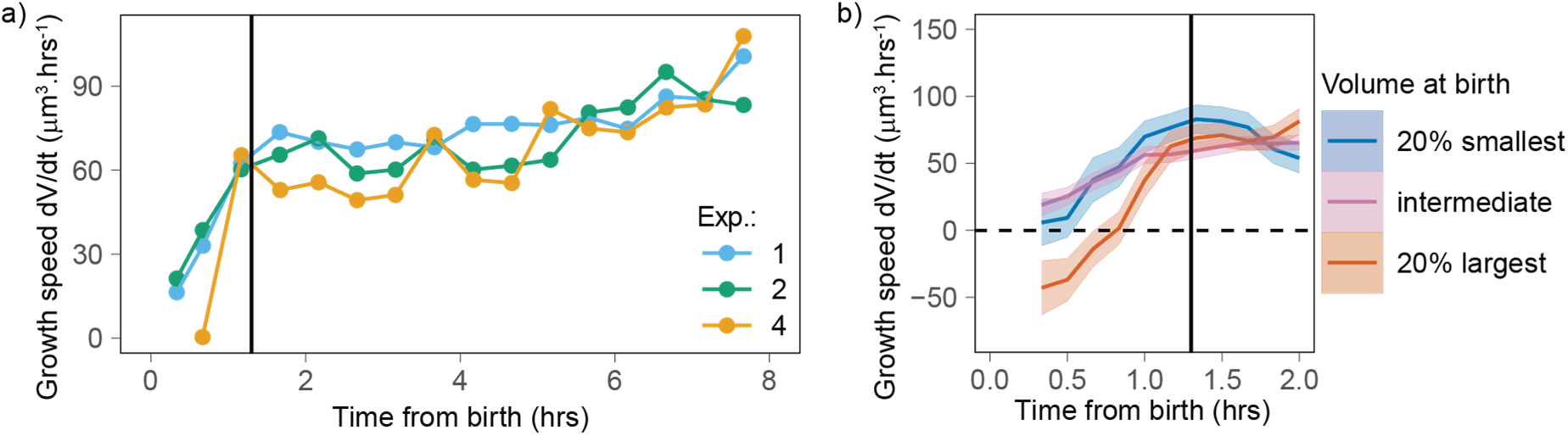
Newborn cells show a different trend in growth speed. a) Growth speed (proxied by the discrete derivative 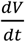 taken on 50 min windows) as a function of time from birth (onset of cytokinesis) for the three experiments that yielded more than 80 different cells. The vertical line indicates 1.3 hours after birth. Circles are averages bins containing at least 100 cells (N=3). b) Zoom of the first 2 hours of the data shown in panel a with the data pooled by volume at birth (blue, orange, and pink lines correspond, respectively, to sliding averages of the largest 20% birth volume, smallest 20% birth volume, and the rest of the cells, the dashed horizontal line indicates a growth speed equal to 0, the black vertical line indicates 1.3 hours from birth).

### Volume growth rate fluctuations decrease with cell volume

Our analyses so far indicate that, excluding the initial (Figure 3) stage of the cell cycle, volume growth is close to exponential for a wide range of volumes (Figure 2a), at a rate that changes with cell-cycle phase (Figure 2c). Next, since little is known about the cell-specific variations around this average behavior, we set out to evaluate the fluctuations, focusing on the range of volumes for which the growth behavior is well characterized (cells between 1800 and 3100 μm^3^ (Figure 2a) and starting 1.3 hrs after birth (Figure 3a)). Volume growth is the result of the combination of both biosynthetic pathways that act over the cell cycle and homeostatic pathways (that typically act at shorter time scales) that maintain a balance of cellular osmosis, hydrostatic pressure, and density^1,31^. We first quantified the variability of volume growth by the variance of the specific growth rate, proxied by 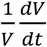, for cells grouped in different volume bins, and we found that this variance decreases rapidly with the time scale *dt* over which one takes the discrete derivative (Figure 4a). Hence, the time scale must be specified for a meaningful comparison of the size of such fluctuations (for example across different studies). Moreover, we found that, at fixed time scale of the discrete derivative, the variance in specific growth rate decreases with volume (Figure 4a). Since this observation was robust across all derivative time scales, we decided to focus on the fluctuations of growth rate measured at the shortest accessible time scale (50 minutes, corresponding to 5 frames). When we plotted together the mean and standard deviation of specific growth rate as a function of volume (Figure 4b), the standard deviation of specific growth rate clearly decreases with volume.

**Fig. 4:**
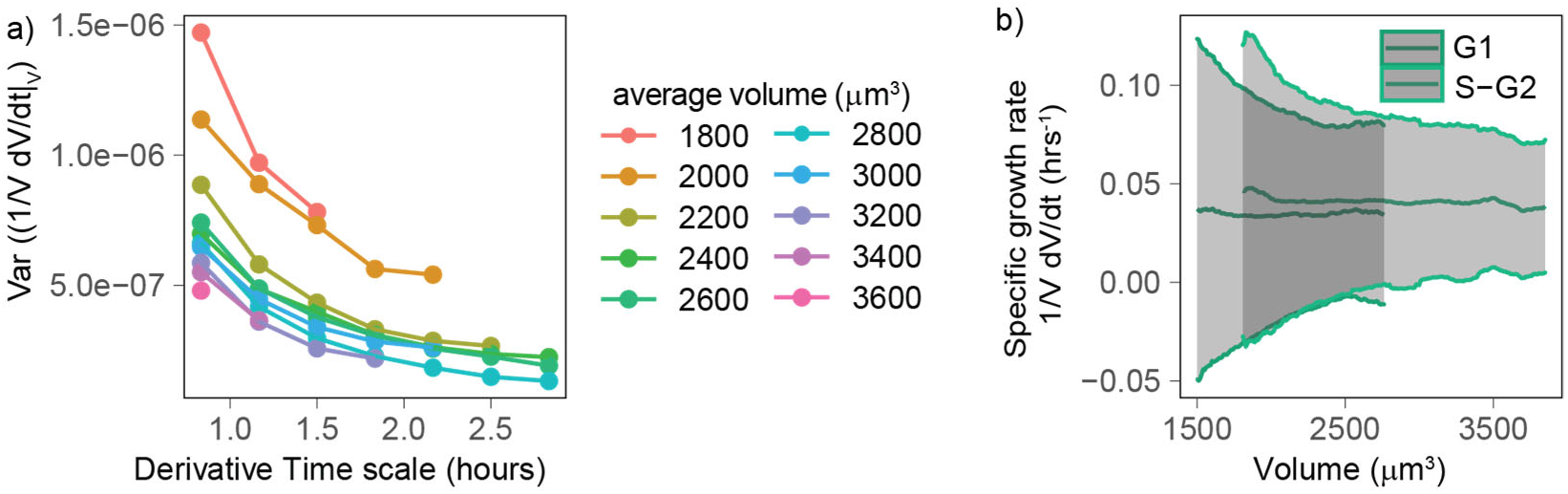
Volume growth rate fluctuations decrease with increasing cell volume. a) Variance of specific growth rate quantified by 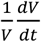 using discrete derivatives of fixed volume bins calculated over increasing time windows (x axis) and for groups cells of increasing volume (colored lines). Circles are averages computed for bins that contain at least 100 different cells, N=4. b) Mean (line) and standard deviation (grey ribbon) of growth rate (quantified by 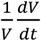 using discrete derivatives on fixed volume bins and a time scale of 50 mins) plotted as a function of volume and by phase. Values are calculated on sliding windows of 200 μm^3^ and bins contain at least 100 cells (N=4).

### Growth rate fluctuations are dominated by constant noise

To better understand the observation of a reduction in growth rate noise with cell volume, we used a stochastic mathematical model describing cell growth (see Annex and Figure 4a). This model has a long history of applications outside biology^32,33^, but it was recently proposed by Pirjol and colleagues^34^ in the context of cell growth. The model considers that cells on average grow exponentially and describes fluctuations around this mean growth as noise:

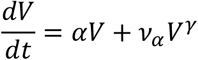

where *α* is the mean specific growth rate and *ν_α_* is a white noise term. Thanks to the factor *V^γ^*, this equation interpolates the limit cases of “additive” (constant amplitude) fluctuations and “multiplicative” (volume-specific) fluctuations (Figure 5a, see also Annex A for details and Figure 5 – supplement 1a for the validation with simulations). Central to the model is the definition of the parameter *γ* (0≤ *γ* ≤1) which sets the relative weight between two kinds of noise: when *γ* = 1, we obtain: *dV/dt* = (*α* + *ν^α^*)*V*, and the model describes multiplicative specific growth rate fluctuations (Figure 5b). These fluctuations are symmetric with respect to a reference specific growth rate *α*, hence they can be interpreted as emerging from noise in biosynthetic rates (e.g., surface synthesis, protein synthesis, etc.). When *γ* = 0, the model describes an additive noise of constant amplitude, acting symmetrically on growth speed *dV/dt*, which can be interpreted as resulting from any of the homeostatic processes that contribute to setting steady state volume (e.g. homeostatic constraints of biophysical origin on hydrostatic pressure, osmotic pressure, etc.). The model also allows for intermediate values of *γ*, which would effectively describe the combined presence of additive and multiplicative noise sources on growth.

**Figure 5:**
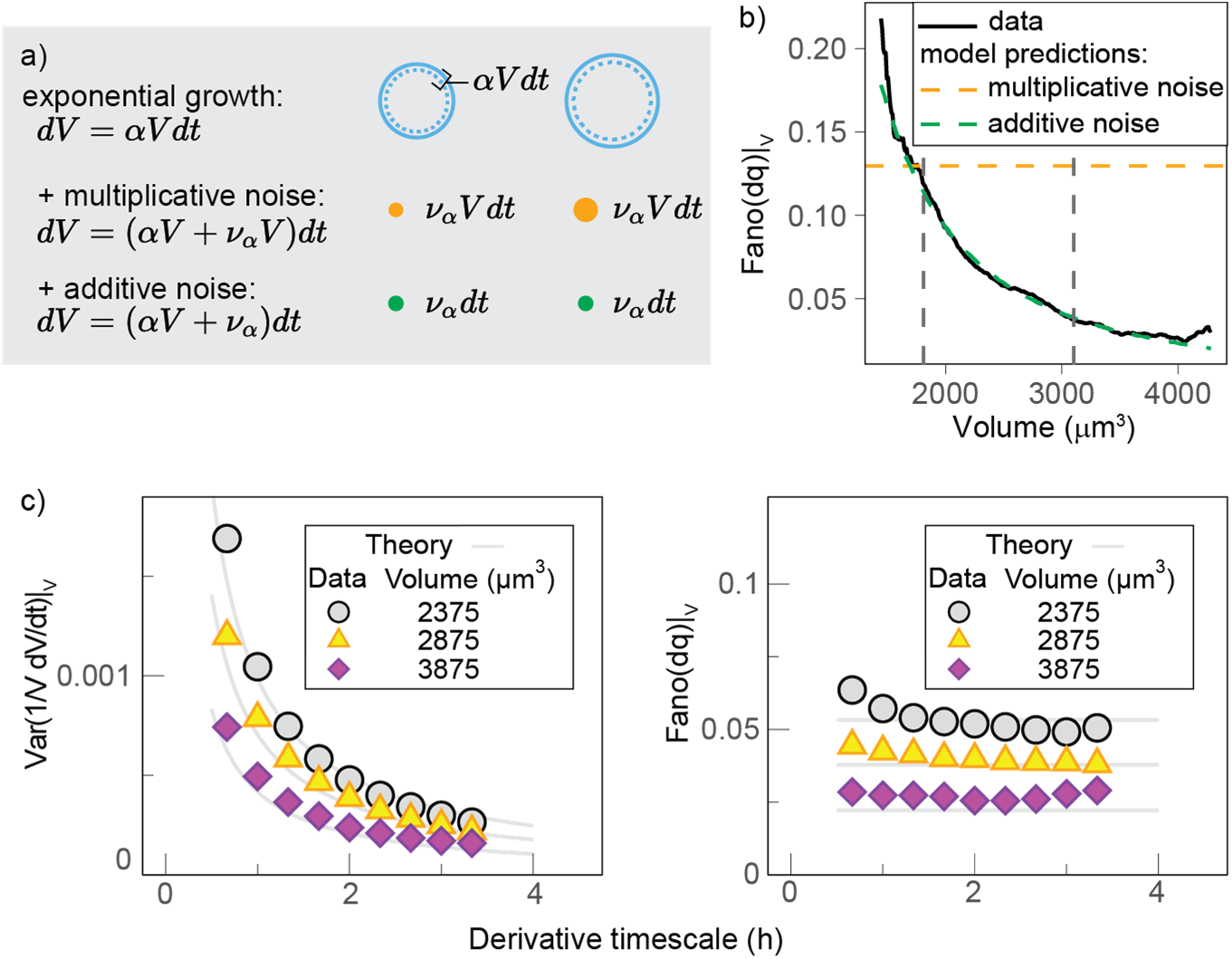
Growth rate fluctuations are best described by a model with additive noise. a) Schematic representing the exponential growth of a cell (dashed blue circle) adding volume *dV* (solid blue circle) proportionately to its volume at a rate *α* and to time scale *dt*. Two limit-cases for fluctuations around this baseline exponential growth are a “multiplicative noise”, which is volume-specific, hence increases with volume, or an “additive” noise, whose amplitude is constant. b) Mean-normalized variance (Fano factor) of the conditional “log return” 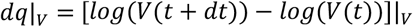, computed in sliding volume bins, and plotted as a function of volume. The dashed lines represent the theoretical prediction of the model in the case of pure multiplicative (orange dashed line) or pure additive noise (green dashed line), assuming that the amplitude of the noise is set by the fluctuations in the smallest volume bin (yielding a constant line with an intercept *b* = *Fano*(*dq*|_*V*=1800μ*m*3_) for multiplicative noise, and a power law ~*b/V*^2^ with the same condition for additive noise). c) The Fano factor quantifies fluctuations robustly over different time scales. Left panel: comparison of the variance specific growth rate (quantified by 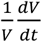 using discrete derivatives on fixed volume) as a function of the time scale of the discrete derivatives in the data (symbols) and the theoretical model predictions (solid grey lines) for different volume bins. Right panel: the Fano factor of the conditional log return *dq*|_*V*_ defined above (same symbols), is robust to time scale changes, as predicted by the model.

An important prediction of the model (Figure 5c) is that the mean-normalized variance of growth rate (Fano factor) is not dependent on the time scale of the derivative. When we examined our experimental data, we observed that this was indeed the case (Figure 5c). The Fano factor of growth rate depends on *γ*, the mean specific growth rate *α* and constant *σ* as follows:

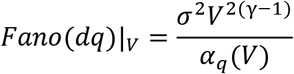

Where *α_q_* is the average log-return rate at fixed volume (see Annex A), and 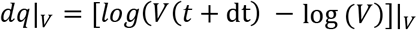 is a conditional “log return” and quantifies the volume specific growth rate in presence of possible multiplicative fluctuations. Importantly, other estimators of variability such as variance, SD or CV, do depend on the time scale of the discrete derivatives, in both the model and the data.

We thus decided to use the Fano factor to estimate the variability in growth rate. We found that the Fano Factor of volume growth rate (measured using the “log return” *dq*) decreased rapidly with volume and that the trends measured in G1 and S-G2 overlapped well (Figure 5 – Supplement 1b). This suggests that the decay in volume growth rate variability is independent of cell cycle phase. We thus pooled the two phases together and compared the decay with our model predictions (Figure 5b). Comparison of the model predictions for the two extreme cases (*γ* = 1 and *γ* = 0) and the data shows that the data are very close to a scenario where *γ* ≃ 0 and fluctuations are almost entirely additive (Figure 5b). Such additive noise could arise from an extrinsic additive noise on volume measurement. To estimate the noise on volume measurement, we compared the fluctuations on the detrended volume curves of cells and neighboring background areas of same surface area (Figure 5 - supplement 2 a-b). The standard deviation of the fluctuations was 3-4 times larger for cells than for the background areas, showing that fluctuations of biological origins dominate over fluctuations of technical noise origin (Figure 5 - supplement 2c). Overall, this analysis suggests that volume fluctuations come principally from a constant-amplitude noise in instantaneously added volume, rather than from random variations of the volume-specific growth rate itself.

## Discussion

Most recent studies on animal cell growth have considered the patterns of dry or buoyant mass^9,35^. Here, we complement this knowledge by providing a high-throughput dataset of volume trajectories in adhering cells allowing us to compare the behavior of these different parameters.

During the first 1.3 hours following birth, volume growth displays a pattern where cells of intermediate and small volume grow slowly while larger cells at birth lose volume. This is very different from the mass pattern previously reported^23^. Our volume curves exclude the period in mitosis where cell volume transiently increase by 10-30% in volume called the mitotic volume overshoot^11,24^. This 1.3 hours period also spans both the period of completion of cytokinesis and the beginning of G1 phase, ruling out the hypothesis that mitosis alone determines the early volume growth pattern. Our observations instead point to the existence of a previously overlooked period early in the cell cycle where several processes such as the change in cell shape due to the spreading of the cell after mitotic rounding^36,37^, and the onset of cell growth in early G1 compete to determining volume growth. It is possible that this period also corresponds to an extreme case where volume and mass growth rate, which are very different during mitosis^11,23,24^, progressively ‘tune in’ before they reach a common steady behavior.

Throughout the cell cycle, mass growth has been reported to oscillate periodically in HeLa cells^9,35^. We find that this phenomenon is not simply reflected by cell volume. We found some oscillations in volume growth speed as a function of time from birth only in birth volume outliers, and not matching the period and amplitude of the mass oscillations reported previously.^9,35^ (Figure 3 – Supplement 1). This finding suggests that dry-mass biosynthesis and volume growth, while being interrelated, can be independent in specific phases of the cell cycle. Studies combining measurements of cell shape, mass, and volume at high time resolution will be particularly important to clarify the complex interplay between these parameters.

The high number of curves in our study allows us to investigate systematically the fluctuations in volume growth. Our observation that volume growth shows additive fluctuations counters the idea that the volume specific growth rate itself fluctuates (Figure 5a-b). What could be the origin of such additive noise and how can we explain the absence of noise on the rate itself? Our controls show that extrinsic experimental noise only minimally contributes to the additive fluctuations we measure (Figure 5 – Supplement 2). Intrinsic sources of additive fluctuation could arise from noise in the processes involved in volume regulation. Cell volume is a physical parameter that results from an equilibrium of cell hydrostatic pressure, osmolarity and membrane tension at time scales of minutes to hours^1^ and much less understood mechanisms that maintain cell density^31,38–41^ by coupling mass and volume at timescales of several hours. In a scenario where volume is unidirectionally coupled to mass, even if mass growth rate is noisy^42^, one could potentially obtain an apparent volume growth rate with fluctuations that are only due to the coupling. None of the available studies considering cell mass has addressed the question of whether mass biosynthesis fluctuations are themselves mass specific.

Finally, our analysis shows that growth-rate variability, quantified by variance, SD, or CV, is strongly dependent on the time scale used to evaluate discrete derivatives (Figure 4a, Figure 5c). This result poses an important caveat for the quantitative comparison of growth rate fluctuations across different studies, as absolute values of growth rate fluctuations evaluated in different ways and on different time scales (or smoothing windows) may strongly differ. In general, single-cell growth studies are currently limited by the development of theoretical tools that could quantify the contribution of the different determinants of growth such as size, time, and cell cycle phase that act at different time scales. These tools, together with experimental approaches that allow the combined measurement of several size parameters (mass, volume) concomitantly are needed to further elucidate the growth patterns of animal cells.

## Materials and Methods

### Cell line and cell culture

HeLa cells expressing hgemin-GFP were a kind gift from Buzz Baum’s lab (UCL, London, United Kingdom). Cells were cultured in DMEM-Glutamax media and imaged in DMEM without phenol red, supplemented with Glutamax. Both media were supplemented with 10% FBS and 1 % penicillin-streptomycin.

### Volume measurement with FXm

The detailed protocol for FXm is described previously^25^ and the design is described in ref.^10^. Briefly, measurement chambers were replicated in PDMS (crosslinker:PDMS, 1:10). To prevent leakage of the fluorescent dextran from the chamber, 4mm high PDMS cubes were stuck on top of the inlets before punching 2mm diameter holes for every inlet. The chamber was then irreversibly bound to the 35 mm diameter glass-bottom fluorodishes® by plasma treatment, coated with fibronectin 50 (μg.mL^-1^), for about 30 min, then rinsed and incubated in phenol-free media overnight. To ensure that cells were in a similar growth phase when starting an experiment, cells were seeded at constant density (1.9×10^4^xcm^-2^) two days prior to the experiment. The day of the experiment, cells were detached by incubating with EDTA for 15-20 min, recovered and seeded at intermediate density in the measurement chamber (Figure 1a). 4 hours after seeding, the media was replaced with equilibrated media containing 1 mg.mL^-1^ of 10 kDa Dextran Alexa-645. Imaging started 2-4 hours after changing the media. While imaging, cells were kept at 37°C with 5% CO2 atmosphere. Imaging was performed on an inverted epifluorescence microscope (Ti inverted (Nikon) or DMi8 inverted (Leica)) equipped with a LED excitation source. Images were acquired with a CoolSnap HQ2 camera (Photometrics) or an ORCA-FLASh4.0 camera (Hammamatsu). Images were obtained using a low magnification (10X), low numerical aperture objective (NA=0.3, phase) every 10 min (FXm measurement) and 30 min (hgeminin-GFP imaging).

### Software analysis

To extract cell volume and cell cycle information from the images we used a custom-made Matlab software^25^. The software contained an image analysis algorithm previously optimized^25^ that performed successive image treatments to normalize the background intensity and correct for background inhomogeneity (for example due to an inhomogeneous light source). The algorithm then segmented the pillars and background to calibrate the fluorescence intensity signal using: (i) the average background intensity to calculate *I_max_*, (ii) the average intensity under the pillars in the chamber to calculate *I_min_* and (iii) the known height of the chamber. The software then segmented and tracked single cells (Figure 1b) throughout the duration of the movie. If a cell divided, the event was recorded and the lineage tree for that cell recorded. Finally, the cell volume and hgeminin-GFP intensity was calculated over the segmented area for each cell and each movie frame.

The background normalization algorithm required the user to manually set several parameters. To ensure that these parameters were accurately chosen, a graphical user interface allowed the user to visualize the results of the image treatment given a set value for each parameter and assess its validity. This allowed, after a few trial and errors, setting a set of parameter values that were optimal for each set of movies obtained in the same FXm chamber. There were four steps to set these parameter values. First, to detect the pillars and later calculate *I_min_*, the user manually set a threshold that segmented the pillars (from 0 to 1 on a normalized image), the user also set a ‘distance of influence’ around each pillar which consisted in an area larger than the pillar where cell volume would not be calculated to prevent potential artefacts of volume calculation due to a shadow caused by the pillars (see ref.^25^). Second, to estimate the background and later calculate *I_max_*, the user chose a threshold to detect the cells (from 0 to 1 on a normalized image) and set the parameter called ‘noise factor’. Third, using these parameter values, the image treatment algorithm was applied to the image. Fourth, on the resulting renormalized image, the user set the parameter values for cell segmentation and tracking: the average size of the mask and the threshold value to detect the cell, the maximum moving distance for a cell from one image to the next, the radius around each detected cell (to prevent measuring cells that were too close to each other), the minimum cell size and a parameter ‘sigma’ that represented a threshold allowing splitting of an object into two distinct object (for example after cell division or when two cells are near each other).

During the optimization phase of the analysis pipeline, we re-ran the analysis on the same movies using different parameter values to test the robustness of the volume calculated to variabilities in user-defined parameter values. The volume curves obtained were very similar and indicated that errors in volume measurement due to variability in parameter settings were negligible. Once the algorithm parameters were set, the software processed hundreds of movies in batch mode using parallel processing to increase the speed of processing. The analysis of a set of movies coming from one experiment took 2 to 3 full days of computer processing. At the end of this step, we obtained hundreds of movies showing the tracking results (Figure 1c). Each movie was then visually checked to correct any errors of segmentation.

### Visual assessment and manual curation of the single cell tracks

Each single-cell trajectory was visually checked. We verified that the segmentation and lineage tracing (recording of mitotic events) were accurate. During this manual curation, we visually assessed and noted, for each cell: (i) potential frames when the volume calculated should be excluded from the analysis due to a segmentation error, (ii) if the cell divided, the frame when the cell started rounding and the frame when the first evidence of cytokinesis occurred. We also checked for and corrected mistakes in the lineage tracking. For the first experiment analyzed, we also noted any frames where a cell was near another cell (typically right after birth when the two daughter cells are near each other or later when two cells bump into each other). We then checked that the presence of a neighboring cell was not affecting the volume curve in an obvious way. Since it did not, we stopped tracking this information for the subsequent experiments. This visual assessment and manual curation, although time consuming, was essential to our analysis because it increased our confidence that any fluctuation seen on the resulting volume curve was not due to an identifiable artefact. The resulting volume curves and hgeminin-GFP signal were then imported into R. Using a graphical user interface, each curve was plotted and the user manually selected, for each cell: (i) the beginning and the end of the mitotic volume overshoot^10,11,24^ on the volume curve and (ii) the point of increase in hgeminin-GFP intensity indicating G1/S transition (Figure 1d).

### Volume curve smoothing and calculation of growth speed

To then get into the analysis of volume growth speed and growth rate fluctuations, we developed a cleaning and algorithm that would filter out the clear outliers and smooth fluctuations that are within the noise of our measurement technique. Several algorithms were tested, each time checking visually the resulting comparison of the raw volume measurement with the smoothed, filtered curve. The final algorithm selected worked in two steps. First, to filter out clear outliers, a histogram of values on sliding windows of 11 frames was established and the fourth quantile (*Q*_4_) and interquartile range (*IQR*) of that distribution were calculated. Then, points that were above or below *Q*_4_ + 0.9 \**IQR* were removed from the volume curves. Second, a smoothing algorithm based on centered averages on sliding windows of 3 frames was applied. To calculate growth speed 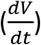, local robust linear fits of single cell volume curves as a function of time were performed on sliding windows of 5 frames (all figures except Figure 4a where the fits were performed on increasingly long windows of time). The slope coefficient of the fit corresponds to the instantaneous growth speed. We compared this approach to calculating the discrete time derivative by plotting the resulting local fits from both methods and visually concluded that the robust linear fit method gave a more faithful representation of the curve fluctuations.

### Model and simulations

The model and analytical calculations to estimate cell-to-cell variability are presented in Annex A. These calculations where compared with simulations based on a discrete-time realization of the Langevin equation defining the model (See Figure 5 - supplement 1).

### Statistical analysis

All figure generation and statistical analysis was performed in R. Packages used were ‘ggplot2’, ‘gridExtra’, and ‘robustbase’.

## Authors contributions

C.C. and M.P. designed the experiments, C.C. and M.C.L. designed the data analysis, C.C. performed experiments and data analysis, M.C.L. and C.C. worked on model and simulations, and performed model-related data analysis. C.C., M.C.L. and M.P. wrote the paper.

## Acknowledgements

Part of this work was carried out at the Aspen Center for Theoretical Physics. The experiments were performed at the Institut Pierre-Gilles de Gennes (IPGG) and the Institut Curie. C.C. would like to thank the imaging platform from the Institut Curie PICT-IBiSA and the IPGG, and James Utterback for advice on the simulations. The authors would like to thank Teemu Miettinen, Ethan Levien, Orso Maria Romano Ludovico Calabrese and Gabriele Micali for helpful conversations, and Matthew Swaffer and Helena Cantwell for comments on the manuscript. MCL was funded by the Italian Association for Cancer Research AIRC-IG (REF: 23258)

## Competing interest

The authors declare no financial and non-financial competing interests.

## Software

Matlab® (MathWorks, Natick, Massachusetts, USA) was used for the image analysis (version R2018a or later).

## Supplementary figures

**Figure 1 – supplement 1:**
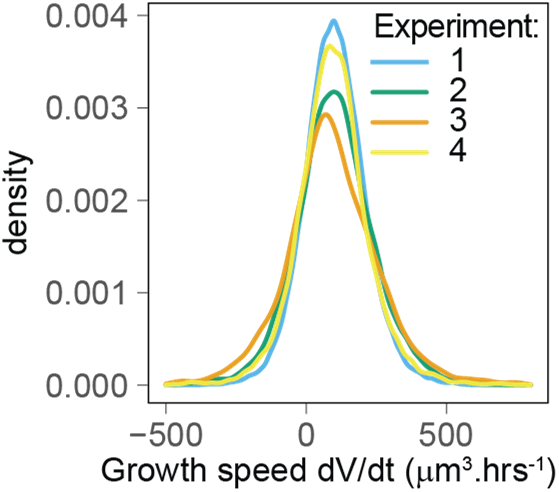
The distributions of growth speed show good agreement between the 4 experimental replicates. Experiment 1: mean±SD=102+/-111 μm^3^.h^-1^, n=50708; experiment 2: mean±SD=102+/-142 μm^3^.h^-1^, n=23739; experiment 3: mean±SD=101+/-196 μm^3^.h^-1^, n=4570; experiment 4: mean±SD=96+/-138 μm^3^.h^-1^, n=12494.

**Figure 3 - supplement 1:**
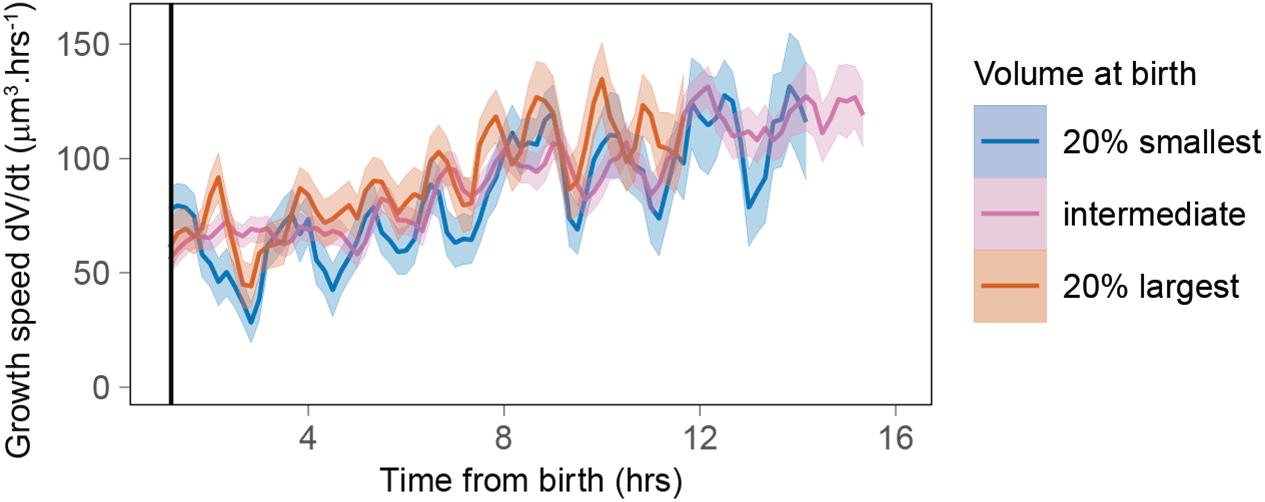
Growth speed as a function of time from birth shows time periodicity for the largest and smallest cells at birth. The solid lines represent averages (each bin contains at least 100 cells) and the ribbon the standard error on the mean. Black vertical line indicates 1.3 hours after birth (N=4).

**Figure 5 – supplement 1:**
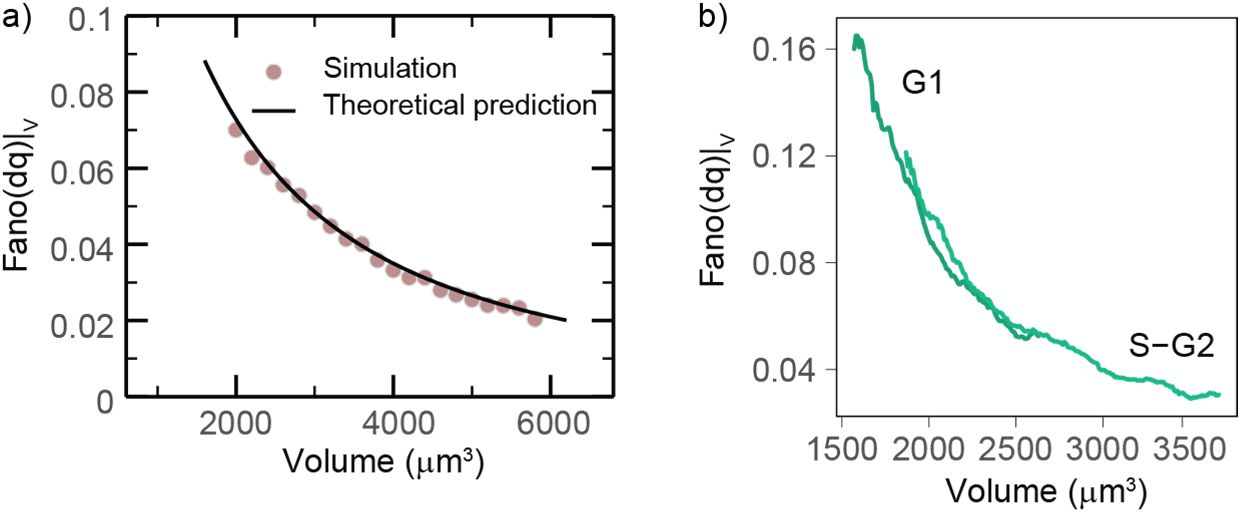
Validation of the model. a) Theoretical model predictions agree with direct simulations of the stochastic exponential growth model. The plot compares simulations (circles) with the theoretical predictions (lines) for the Fano factor of the conditional log return (the logarithmic size increment at fixed volume) from the same set of parameters. Model parameters were derived from a fit of the pulled experimental data from experiments 1 and 2 in the volume range 1800-4000 μm^3^: α=0.038.h^-1^, σ = 41.5(μm^3^)^1-γ^.h^-(1/2)^, γ=0.12. b) Mean-normalized variance (Fano factor) of the conditional “log return” 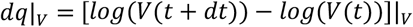, computed in sliding volume bins, and plotted as a function of volume and phase.

**Figure 5 – supplement 2:**
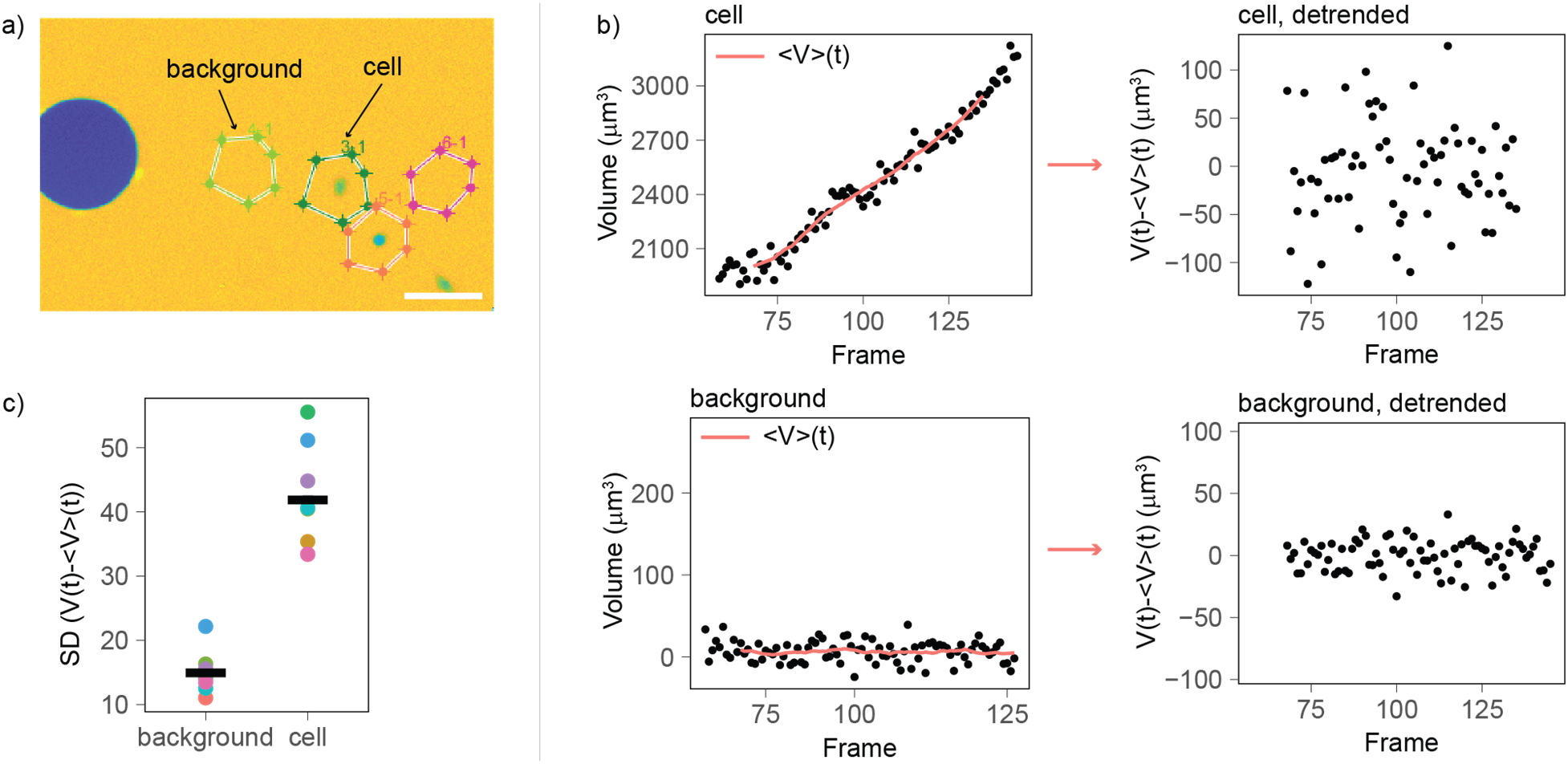
Comparison of the volume fluctuations on cells (biological fluctuations) and background areas (measurement noise). a) Image of the volume analysis software showing two cells and two background areas of same surface area (scale bar: 100 μm). b) Left: For each cell and background area, volume was calculated over time, and an average volume trend computed over sliding windows of 10 frames was calculated (solid red line). Right: the average volume trend was subtracted from the volume curve to obtain a detrended volume curve. c) To estimate the volume fluctuations around the mean detrended curves, the standard deviation was calculated for each cell (6 total) and corresponding background areas (colors of the dots correspond to a pair of cell and matching background area). The volume fluctuations measured for cells are about 3-4 times higher than those measured on background areas, suggesting that fluctuations of biological noise origin dominate of fluctuations of technical noise origin.

## Annex A: Stochastic model of exponential growth with noise

This annex describes the stochastic growth model developed to interpret the reduction of noise on growth rate as a function of volume (Figure 5). We compared our data with the stochastic growth model of ref. 9. Following this study, we write

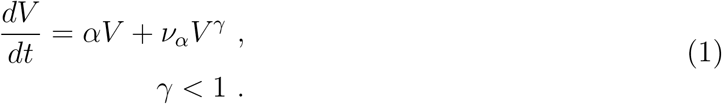

This model interpolates the regimes of additive noise (*γ* = 0) from fluctuations of biophysical nature and corresponds to purely multiplicative noise (*γ* = 1), corresponding to constant-amplitude fluctuations of the specific growth rate around its typical value *α*. The intermediate case *γ* ∈ [0,1] effectively describes a combination of biophysical noise and growth rate fluctuations Here we assume that *ν_α_* is a zero-mean delta-correlated Gaussian white noise, i.e. 〈*ν_α_*(*t*)〉 = 0, and 〈*ν_α_*(*t*)*ν_α_*(*t* + *τ*)〉 = *σ*^2^*δ*(*τ*).

This model was used to compute analytically different proxies of growth-rate fluctuations, which we used to describe the data. We also verified the agreement of our theoretical predictions with direct numerical simulations of Eq. (1).

For *γ* < 1. Let us look at the expected rate and the rate of the mean in this case. Averaging Eq. (1) one has that

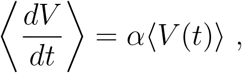

because the average of the noise is zero. Hence we always have that

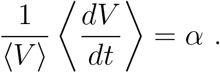

The growth rate in the model can be estimated as the conditional average of the speed at fixed volume. Indeed, since

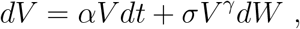

One immediately has that

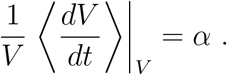

Considering the variance, with some straightfoward algebra one can obtain from Eq. (1) the expression

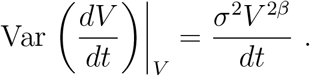

This expression is compared to data in Figure 5c. Defining *q* = log(V), one has that

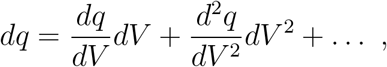

hence

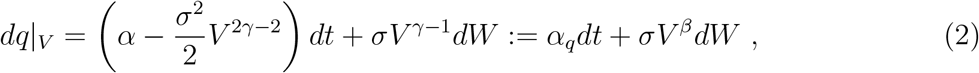

where we have defined *β* = *γ* — 1 and

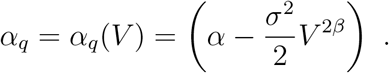

From Eq. (2), we compute with some algebra the Fano factor of *dq* (variance over mean), conditional to volume, which is not dependent on the time scale *dt*,

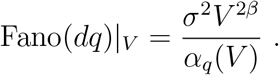

This expression is compared to data in Figure 5c.

## Notes

### Competing Interest Statement

The authors have declared no competing interest.

